## Supplementary Figures for "Spectral Hallmark of Auditory-Tactile Interactions in the Mouse Somatosensory Cortex"

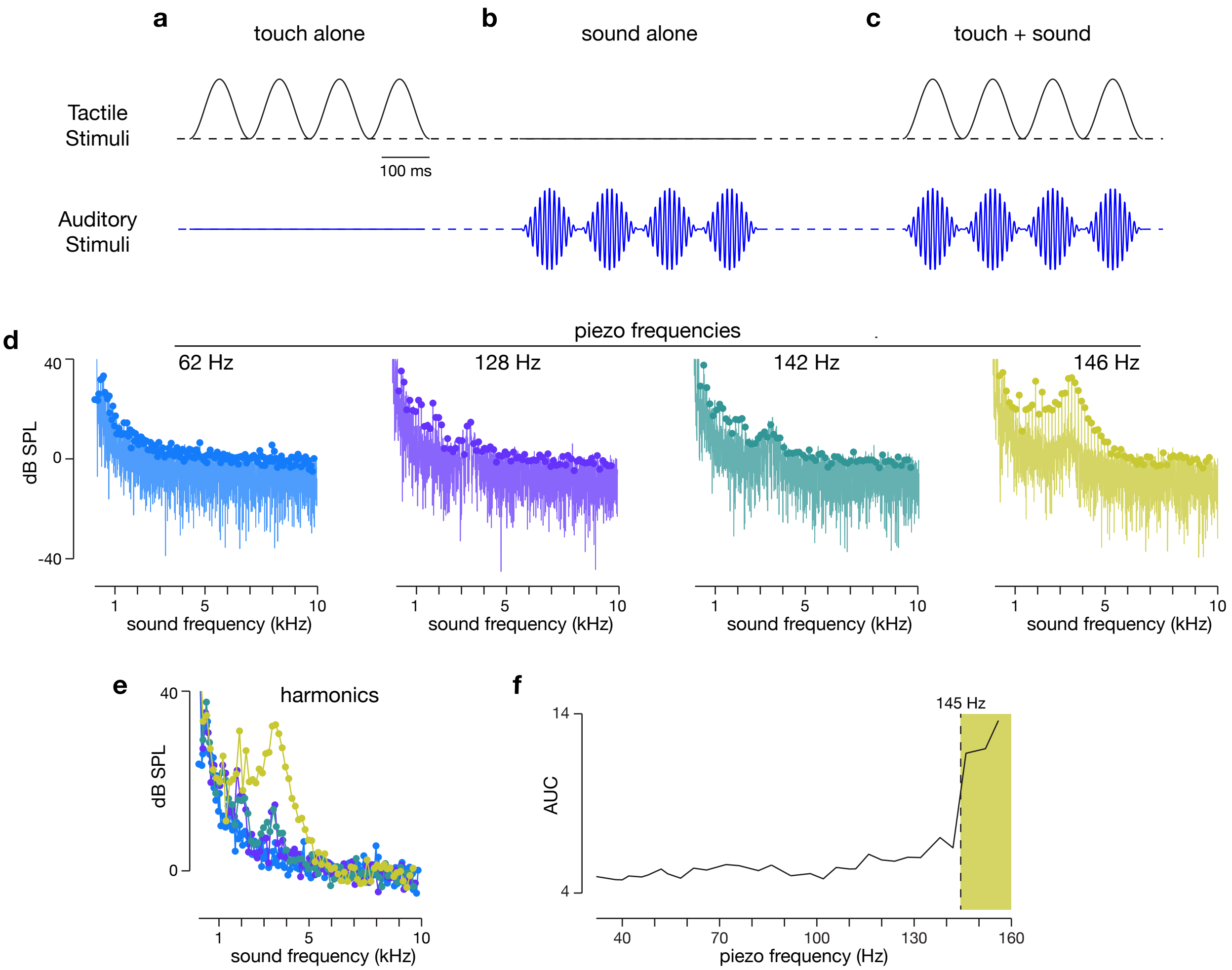


**Supplementary Figure 1 | Schematic Waveforms for Three Different Stimuli. a** Tactile stimulation alone, illustrated by a waveform of an 8 Hz sinusoidal tactile stimulus with a 500 ms stimulus period. 100 ms scale bar shown at the bottom right. **b** Sound stimulation alone, illustrated by a waveform of a 100 Hz tone with an 8 Hz SAM envelope, again with a 500 ms stimulus period. **c** Combined auditory-tactile stimulation, with simultaneous presentation of tactile (top) and auditory (bottom) stimuli. Here the 8 Hz frequency of the tactile stimulus is matched with the frequency of the SAM envelope of the sound stimulus. **d** Spectrum of sound generated by piezo at 62, 128, 142, and 146 Hz (left to right) as measured by probe tube microphone. Local maxima highlighted by dots along the spectrum and are observed at integer multiples of the frequency used to drive the piezo. **e** Values of harmonic peaks are plotted on the same axis. Colors correspond to colors used in **d**. **f** The total area under the curve (AUC) for harmonics at each piezo frequency tested (32 to 156 Hz). Shaded region illustrates the frequencies at which the piezo generated increased sound amplitudes.

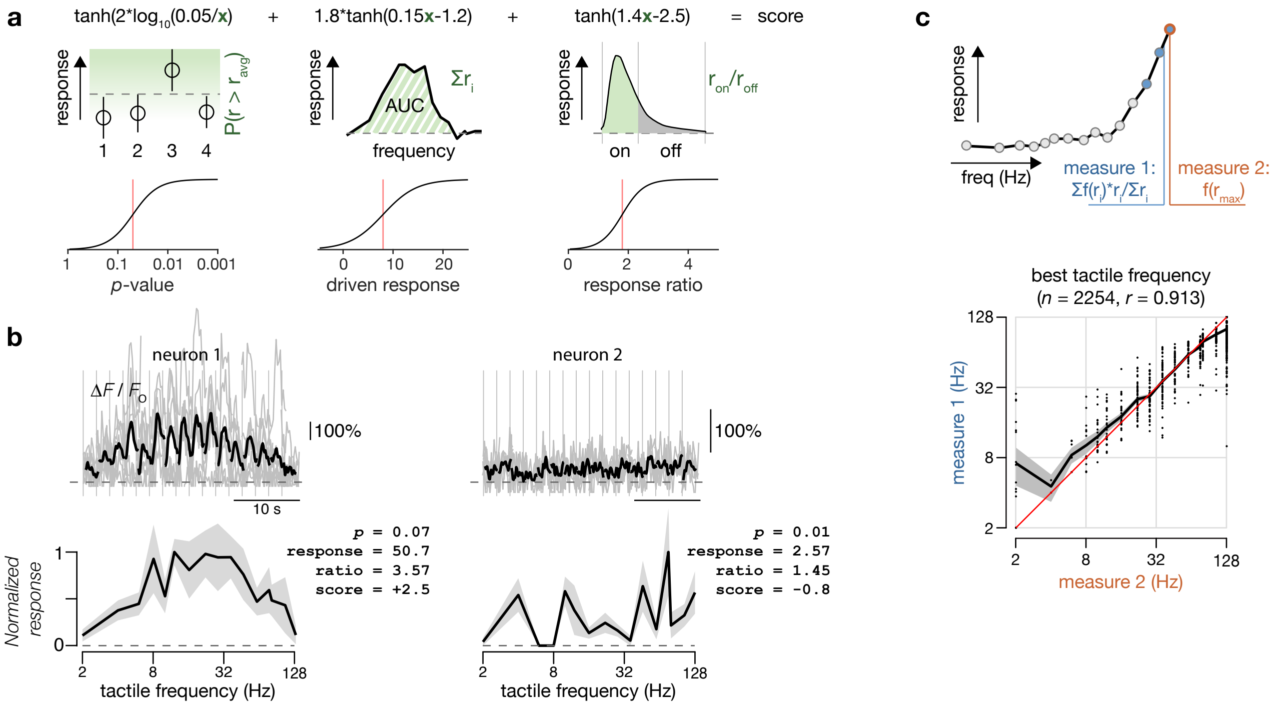


**Supplementary Figure 2 | Measures of stimulus responsivity and best tactile frequency.**
**a** Response score was summed across three measures (*p*-value from 2-way ANOVA, area under the curve of the driven response, and ratio of driven response to spontaneous activity) after passing through a hyperbolic tangent function. Scores ranged from -3.5 to +3.8 with values greater than 1.0 deemed as significant. **b** Tuning curves and responsivity scores for a responsive (left) and an unresponsive (right) neuron. Top panel: individual (gray) and mean (black) responses to individual stimuli. Bottom panel: average response (black) with SEM (gray) across tactile frequencies. **c** Two measures were used to assess the best frequency of a response (upper panel), either the weighted sum of the 3 frequencies eliciting the largest response (measure 1, blue) or the single frequency eliciting the largest response (measure 2, orange). The two measures were highly correlated across the population of responsive neurons (bottom panel).

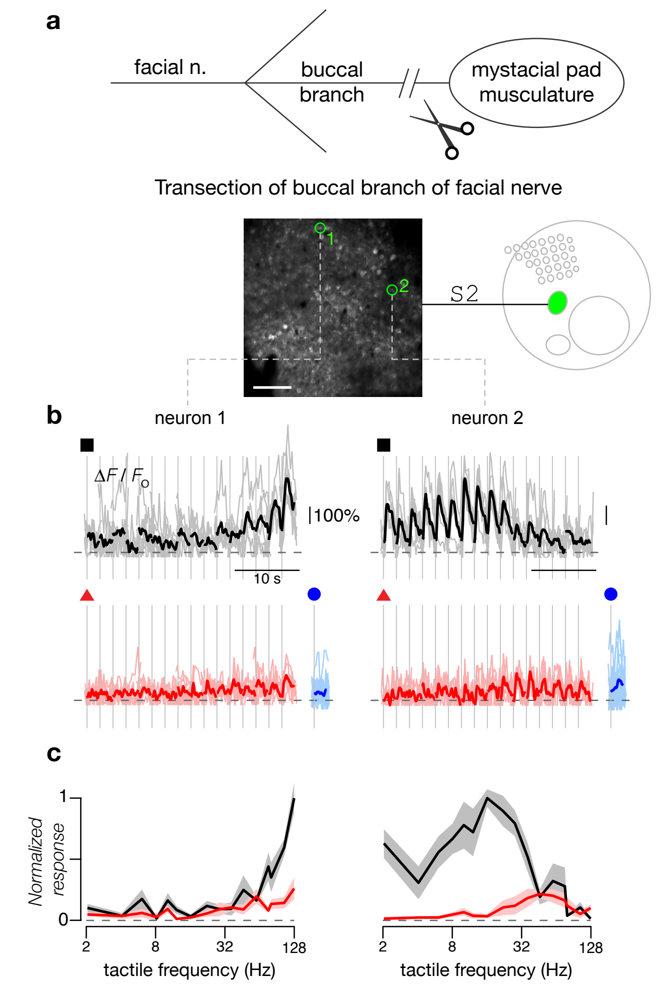


**Supplementary Figure 3 | Preservation of Tactile-Auditory Interactions in the Absence of Active Whisking. a** The buccal branch of the facial nerve was transected in a subset (*n* = 2) of mice to provide temporary (~2 weeks) unilateral abatement of active whisking. Baseline fluorescence image of two-photon imaging field (bottom) located in S2 with exemplar neurons highlighted in green. Scale bar: 100 µm. **b** Exemplar neurons observed in S2 after transection of the facial nerve. Neuronal responses to tactile stimuli alone (black traces) were significantly suppressed when auditory stimuli (8 kHz tones with 10 Hz SAM envelope at 20 dB attenuation) were presented at the same time (red traces). Neither neuron responded to sound stimuli alone (6, 7.5, and 9.5 kHz tones with 10 Hz SAM envelope at 20 dB attenuation; blue traces). Averages are each over 6 repeats. **c** Frequency tuning curves of the normalized response to tactile stimuli and combined tactile plus auditory stimuli for exemplar neurons in **b**. **d** Fraction of neurons in S2 exhibiting either no change, a decrease (inhibition), or an increase (facilitation) when SAM tones were added to the tactile stimuli. Left panel shows data for FOVs imaged in S2 before the facial nerve was transected while the right panel shows data for FOVs imaged in S2 after the facial nerve was transected. Criteria determined by 2-way ANOVA ($p < 0.05$). Data taken from 7 FOVs across 2 animals (94 tactile responsive neurons) before transection and 6 FOVs across the same 2 animals (119 tactile responsive neurons) after transection. **e** Percentage of change in response of S2 neurons to tactile stimuli ($\Delta R$) when SAM tones were added. Neurons were divided by nerve intact (left) and nerve cut (right) conditions that showed either a decrease (gray) or increase (black) in their response. Error bars show the standard error of $\Delta R$ among inhibited or facilitated neurons.

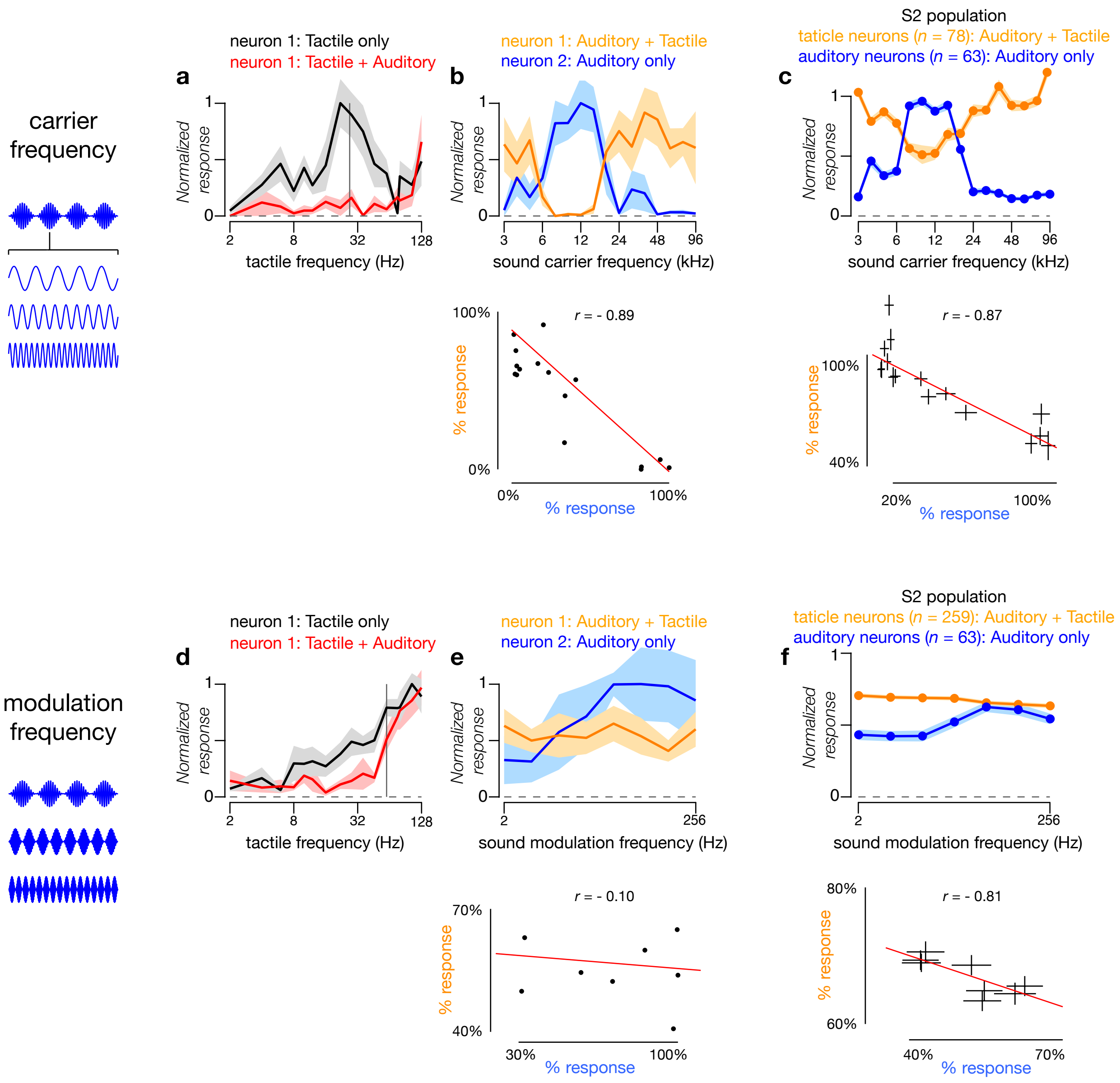


**Supplementary Figure 4 | Independence of Sound-Driven Inhibition to Sound Properties.** Sound-driven inhibition of tactile responses in S2 were not dependent on carrier frequency (**a**-**c**) or modulation frequency (**d**-**f**) of the SAM tone. All error bars are ±SEM. **a** Exemplar tactile-responsive neuron inhibited by concurrent sound. Tactile responses (black curve, 10 repeats), which peaked for frequencies around 20-30 Hz, were mostly abolished by presentation of concurrent sound (red curve, 6 repeats, 12 kHz tone with 10 Hz SAM envelope at 20 dB attenuation). Vertical gray line indicates 28 Hz tactile stimulus to be used in **b**. **b** Change in response as carrier frequency of SAM tone is swept from 3 kHz to 96 kHz. Response of exemplar neuron 1 (same as **a**) to SAM tone during concurrent presentation of 28 Hz tactile stimulus (upper row, orange curve, 7 repeats) shows inhibition of tactile response for sound carrier frequency around 10 kHz. Meanwhile, response of exemplar neuron 2 from the same FOV to SAM tone alone as a function of sound carrier frequency shows the strongest responses around 10 kHz (upper row, blue curve, 6 repeats). Responses of the two exemplar neurons are anti-correlated (bottom row, $p < 0.05$ for Pearson's correlation). **c** For the population of S2 neurons (26 FOVs from 6 mice), sound-selective neurons respond most strongly to SAM tones with carrier frequencies between 6 to 24 kHz (upper row, blue curve), while touch-selective neurons are most inhibited by concurrent SAM tones with carrier frequencies in the same range (upper row, orange curve). This anti-correlation is confirmed by plotting the two responses against each other, parameterized by the sound carrier frequency (bottom row, $p < 0.05$ for Pearson's correlation). **d** Response of exemplar neurons to tactile stimuli without (black, 9 repeats) or with (red, 7 repeats) concurrent sound stimulation (10 kHz tone with 64 Hz SAM modulation frequency at 40 dB attenuation). Vertical gray line indicates 60 Hz tactile stimulus to be used in **e**. **e** Response of exemplar neuron from **d** to SAM tone as a function of modulation frequency during concurrent presentation of 60 Hz tactile stimulus (upper row, orange curve, 5 repeats) and for exemplar sound-selective neuron from the same FOV in response to SAM tones alone as a function of modulation frequency (upper row, blue curve, 6 repeats). The tactile-selective neuron's response is independent of sound modulation frequency ($p > 0.5$, 1-way ANOVA for modulation frequency) and not correlated with neuron 2's response (bottom row, $p > 0.5$ for Pearson's correlation). **f** Average responses of tactile-selective neurons (upper row, orange curve) to SAM tones with concurrent tactile stimulus and of sound-selective neurons (upper row, blue curve) to SAM tones alone, both as a function of sound modulation frequency (43 FOVs from 17 mice). Of the 259 tactile neurons, 20 had a significant dependence on sound modulation frequency (1-way ANOVA with pre-established criterion of $p < 0.05$) while 33 of 63 sound-selective neurons depended on sound modulation frequency. Across all neurons, responses of the two populations were slightly but significantly anti-correlated (bottom row, $p < 0.05$ by Pearson's correlation).

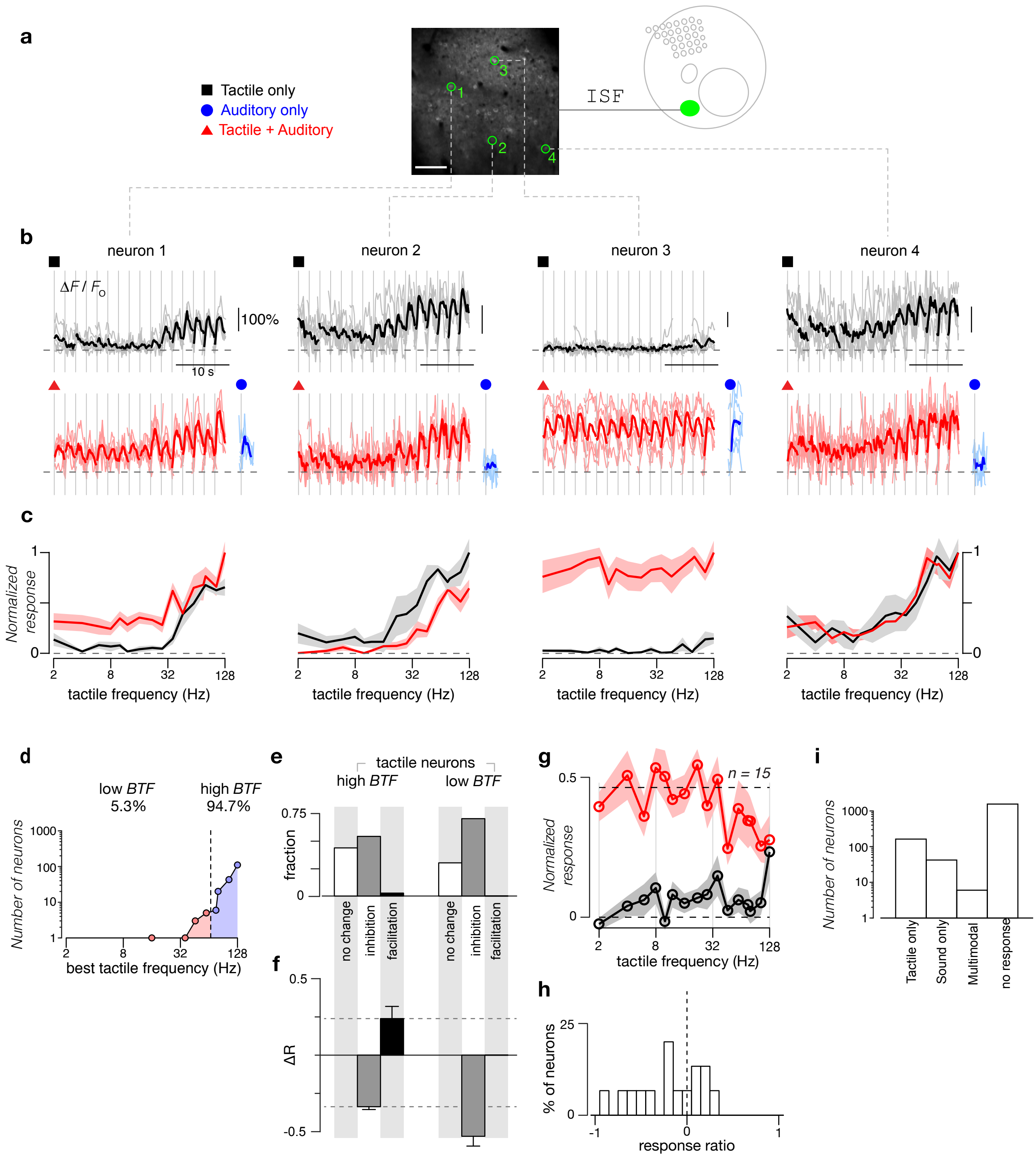


**Supplementary Figure 5 | Diversity of Multimodal Interactions in ISF. a** Imaging FOV located in ISF. Baseline fluorescence image with exemplar neurons highlighted in green. Scale bar: 100 µm. **b** Responses of exemplar neurons to tactile stimuli alone (black traces, top panel) or to combined tactile and auditory stimuli (red traces, bottom panel). For the combination stimulus, a concurrent auditory stimulus (10 kHz tones at 40 dB attenuation) was added to the tactile stimuli. Blue traces (bottom right) show averaged responses to 10 kHz SAM tones with 2 Hz SAM envelope at 40 dB attenuation. Each stimulus was repeated 5 times. **c** Frequency tuning curves of the normalized response to tactile stimuli and combined tactile plus auditory stimuli for exemplar neurons shown in **b**. **d** Population distribution of best tactile frequencies (*BTF,* see Methods) for ISF neurons. Black circles show number of neurons tuned to each frequency tested. Neurons with best frequencies no higher than 60 Hz were categorized as low *BTF* neurons (pink) while those with best frequencies above 60 Hz were categorized as high *BTF* neurons (blue). Both *x* and *y* axes plotted on a logarithmic scale. Data taken from 9 FOVs across 4 mice. **e** Percentage of neurons in ISF exhibiting either no change, a decrease (inhibition), or an increase (facilitation) when SAM tones were added to the tactile stimuli. Criteria determined by 2-way ANOVA ($p < 0.05$). Neurons were categorized based on their *BTF*. **f** Percentage of change in response of ISF neurons to tactile stimuli ($\Delta R$) when SAM tones were added. Neurons were divided into high *BTF* (left) and low *BTF* (right) neurons that showed either a decrease (gray) or increase (black) in their response, as indicated in panel **e**. Error bars show the standard error of $\Delta R$ among inhibited or facilitated neurons. **g** Averaged frequency tuning curves of the normalized response to tactile stimuli and combined tactile plus auditory stimuli for sound selective neurons in ISF that respond to the auditory stimuli used in the combined stimuli ($n$ = 15). Baseline response was additionally subtracted. Standard error shown as shaded gray and red regions. **h** Distribution of response ratio at high frequency relative to low frequency for sound selective neurons in ISF shown in **g**. Response ratio is calculated as $(r_{lo}-r_{hi})/(r_{lo}+r_{hi})$, where $r_{lo}$ is the response to sounds paired with low frequency (2-8 Hz) tactile stimuli and $r_{hi}$ the response to sounds paired with high frequency (76-128 Hz) tactile stimuli. **i** Counts of ISF neurons categorized by response type.
